# Massive rhizobial genomic variations associated with partner quality in *Lotus–Mesorhizobium* symbiosis

**DOI:** 10.1101/2020.03.08.983007

**Authors:** Masaru Bamba, Seishiro Aoki, Tadashi Kajita, Hiroaki Setoguchi, Yasuyuki Watano, Shusei Sato, Takashi Tsuchimatsu

## Abstract

In diverse mutualistic relationships, genetic variations in impact on the growth of interacting partners—variations in partner quality—are common, despite the theoretical prediction that selection favoring high-quality partners should eliminate such variations. Here, we investigated how variations in partner quality could be maintained in the nitrogen-fixing mutualism between *Lotus japonicus* and *Mesorhizobium* bacteria. We reconstructed *de novo* assembled full-genome sequences from nine rhizobial symbionts, finding massive variations in the core genome and the contrastingly similar symbiotic islands, indicating recent horizontal gene transfer (HGT) of the symbiosis islands into diverse *Mesorhizobium* lineages. A cross-inoculation experiment using nine sequenced rhizobial symbionts and 15 *L. japonicus* accessions revealed extensive quality variations represented by plant growth phenotypes, including genotype-by-genotype interactions. Quality variations were not associated with the presence/absence variations of known symbiosis-related genes in the symbiosis island, but rather, showed significant correlations with the core genome variations, supported by SNP- and kinship matrix-based association analyses. These findings highlight the novel role of HGT of symbiosis islands, which indirectly supply mutations of core genomes into *L. japonicus*-associated bacteria, thereby contributing to the maintenance of variations in partner quality.

## INTRODUCTION

While mutualistic symbiotic relationships—unlike organisms living together and establishing a cooperative interaction—are ubiquitous in nature, it remains unclear how mutualistic interactions originated and how they are evolutionarily maintained.

Evolutionary theory predicts that mutualism might be unstable because natural selection would favor mutualists that optimize their fitness by minimizing the costs of returning benefits to a partner (Frederickson 2013; Ghoul *et al*. 2013). Therefore, the stabilizing mechanisms that prevent the invasion of low-quality partners have been a major focus of studies (Heath and Stinchcombe 2014), such as those involving partner choice (Bull and Rice 1991), partner fidelity feedback (Bull and Rice 1991; Archetti *et al*. 2011) and sanctions against such invasion (Kiers *et al*. 2003).

Although such stabilizing mechanisms should reduce variations in partner quality in mutualistic symbiosis, thereby maintaining high-quality mutualists, variations in partner quality have been observed in diverse mutualistic relationships (Thrall *et al*. 2000; Sachs *et al*. 2010;). Multiple models have been proposed to resolve this discrepancy, such as mutation–selection balance and spatiotemporally varying selection (Van Dyken *et al*. 2011; Simonsen and Stinchcombe 2014; Smith *et al*. 2014; Steidinger and Bever 2014; reviewed by Heath and Stinchcombe 2014). The mutation–selection balance is a model postulating that low-quality partners evolve via mutations but are slowly purged from populations by purifying selection. The model of spatiotemporally varying selection predicts that the fitness of a partner’s genotype varies depending on the genotype of an interacting species, such as genotype × genotype (G × G) interactions, on the spatially variable environment conditions (G × E interactions) or on temporally variable selection (Denison and Kiers 2004). While these models provide possible explanations for the persistence of variations in partner quality in mutualist relationships, it remains unclear how variations in partner quality arise and how they are maintained in natural populations.

The legume–rhizobia mutualism is an ideal model to address this question because we can manipulate genotypes of both species and reconstruct their interactions *in vitro*. To understand the origin and the maintenance of variations in partner quality at the microevolutionary scale, it is essential to quantify the quality variations among strains originated from single leguminous species and disentangle the genetic basis underlying the rhizobial quality. Although variations in rhizobial quality have been observed in multiple systems of legume–rhizobia mutualisms (e.g. *Acasia*–*Ensifer*, Barrett *et al*. 2015 and 2016; *Medicago*–*Ensifer*, Porter *et al*. 2011), most previous studies were based on a limited number of genes, and thus, the genetic basis underlying the rhizobial quality was unclear. Although Porter *et al*. (2019) and Klinger *et al*. (2016) analyzed the genetic basis of the rhizobial quality variation using genome-wide polymorphism data, these studies used a limited number of plant strains, and therefore, did not take G × G interactions into account. Furthermore, genome-wide polymorphism data of these previous studies were based on resequencing using Illumina short-reads, making it difficult to investigate structural variations suggested to be important for plant–microbe interactions (Raffaele *et al*. 2010; Tsushima *et al*. 2019).

Here, we focused on the mutualism between *Lotus japonicus* (Regel) K. Larsen and their rhizobial symbionts. *Lotus japonicus* has been regarded as a model species for the understanding of plant–microbe interactions (Bamba *et al*. 2019a), and there have been extensive studies on the molecular, physiological and genomic bases of plant– rhizobia symbiosis (Handberg and Stougaard 1992; Szczyglowski *et al*. 1998; Kouchi *et al*. 2004; Maekawa *et al*. 2009; Madsen *et al*. 2010; Suzuki *et al*. 2011; Soyano *et al*. 2013; Nishida *et al*. 2016, 2018). Among *L. japonicus*-associated symbionts, full-genome sequence information is available from two strains, *Mesorhizobium japonicum* MAFF303099 and *M*. *loti* TONO (Kaneko *et al*. 2000; Shimoda *et al*. 2016). Bamba *et al*. (2019b) explored the genetic diversity of *L. japonicus*-associated symbionts, finding that *L. japonicus* in natural populations were associated with highly diverse *Mesorhizobium* bacteria. However, they used only three housekeeping and five symbiotic genes, and the detailed genomic variations of *L. japonicus*-associated symbionts are still unknown. In this study, to investigate the genomic variations of *L. japonicus*-associated symbionts, we first reconstructed high quality *de novo* assembled genome sequences from nine rhizobial symbionts sampled from three geographically distinct locations in Japan. Second, to quantify the rhizobial variations in partner quality including G × G interactions, we performed a cross-inoculation experiment using nine full-genome sequenced rhizobial symbionts and 15 *L. japonicus* natural accessions.

Three of those *L. japonicus* accessions originated from the same locations where the nine rhizobial strains were collected, so we could explore a signature of local adaptation: i.e. native rhizobial genotypes outperform foreign rhizobial genotypes when associated with the host genotypes originating from the same locations. Finally, to infer which genomic regions were responsible for rhizobial variations in partner quality, we performed a series of analyses testing the association between genomic polymorphisms and rhizobial variations in partner quality.

## MATERIALS AND METHODS

### Bacterial strains

We used nine *L. japonicu*s-associated *Mesorhizobium* strains for this study, which were previously referred to as 113-1-1, 113-3-3, 113-3-9, 131-2-1, 131-2-5, 131-3-5, L-2-11, L-8-3 and L-8-10 (Bamba *et al*. 2019b). These nine strains were sampled from three geographically distinct localities, Tottori (113-1-1, 113-3-3 and 113-3-9), Aomori (131-2-1, 131-2-5 and 131-3-5) and Miyakojima (L-2-11, L-8-3 and L-8-10) (Supporting Information Table S1), where *L. japonicus* natural accessions of the Natural BioResource Project also originated (MG50 from Tottori, MG23 from Aomori and MG20 from Miyakojima; Supporting Information Table S2). Details of the sampling localities have been described by Bamba *et al*. (2019b).

**Table 1.** ANOVAs for growth and nodulation of plants in the inoculum treatments.

|  | d.f. | SDW<sup>a</sup> | NON<sup>b</sup> | NOA<sup>c</sup> | NOAi<sup>d</sup> |
| --- | --- | --- | --- | --- | --- |
| Host | 14 | 0.244*** | 0.228*** | 0.268*** | 0.200*** |
| Inoculant | 8 | 0.093*** | 0.048*** | 0.038*** | 0.081*** |
| Host x Inoculant | 112 | 0.161*** | 0.168*** | 0.145*** | 0.162*** |

<sup>a</sup> Shoot dry weight, <sup>b</sup> Number of nodules, <sup>c</sup> Total size of nodules, and <sup>d</sup> Nodule size per one nodule.
Numerators indicate degree of freedom for d.f and partial $\eta^2$ for each phenotype
An asterisk (\*\*\*) indicates significance at $P < 0.001$

**Table 2.** Mantel test between partner quality variations and kinships of rhizobial genomes

|  | Core<sup>a</sup> |  | Sym<sup>b</sup> |  |
| --- | --- | --- | --- | --- |
|  | Coefficient<sup>a</sup> | P-value<sup>b</sup> | Coefficient<sup>a</sup> | P-value<sup>b</sup> |
| SDW | 0.6561 | 0.0240 | 0.1788 | 1.0000 |
| NON | 0.6556 | 0.0560 | -0.0821 | 1.0000 |
| NOA | 0.1030 | 1.0000 | 0.0729 | 1.0000 |
| NOAi | 0.9333 | 0.0008 | 0.0361 | 1.0000 |

<sup>a</sup>Numerators indicate Spearman's rank correlation coefficient.
<sup>b</sup>Numerators indicate P-value after the Bonferroni correction

### DNA extraction and whole-genome sequencing using MinION and Illumina HiSeq

Prior to DNA extraction, we cultured rhizobial strains on a tryptone yeast (TY) agar plate for 4 days at 28°C, and then picked single colonies and incubated them for 3 days at 28°C in liquid TY medium. After incubation, we precipitated the cells by centrifugation at 13,000 *g* for 3 min and rinsed them with sterilized MilliQ water (Millipore Corp., Burlington, MA, USA) twice. The genomic DNA of each rhizobial strain was extracted using a NucleoBond CB20 system (MACHERY-NAGEL GmbH & Co. KG, Düren, Germany), according to the manufacturer’s instructions. The quality of genomic DNA was confirmed using agarose gel electrophoresis and a BioSpec-nano system (Shimadzu, Kyoto, Japan).

We performed whole-genome sequencing analyses using Oxford Nanopore Technologies (ONT) MinION and Illumina Hiseq 2500 systems. The library for ONT MinION was prepared using Rapid Barcoding kits (SQK-RBK004). We adjusted all nine libraries to the same concentration, mixed them together and then loaded them onto R9.4 flow cells. The sequencing run was performed twice on a MinION MK1b device following the NC_48h_Sequencing_Run_FLO-MIN106_SQK-RBK004 protocol. The library preparation for Illumina HiSeq and sequencing run were performed by DNAFORM (RIKEN, Yokohama, Japan).

### Preprocessing of next-generation sequencing data, *de novo* assembly and annotation

The ONT reads were demultiplexed with Albacore 2.2.2 (https://github.com/Albacore/albacore), and adapter sequences were trimmed with Porechop 0.2.3 (https://github.com/rrwick/Porechop). The quality of demultiplexed reads was calculated with NanoStat 1.1.0 (De Coster *et al*. 2018). The ONT reads were processed with NanoFilt 2.2.0 (De Coster *et al*. 2018) to keep sequences with a q-score > 8, and the first 100 bases were removed to increase sequence quality, with a minimum sequence length of 1 kb.

The overall quality of the HiSeq reads was evaluated using FastQC (http://www.bioinformatics.babraham.ac.uk/projects/fastqc/). After confirming the lack of technical errors in the sequencing, low-quality tails were trimmed from each read using SolexaQA (Cox *et al*. 2010) with a cutoff threshold set at a q-score of 30, and reads shorter than 75 bp were filtered with PRINSEQ++ (Schmieder and Edwards, 2011). The Hiseq-filtered reads with pair-end relationships were repaired with BBtools (https://sourceforge.net/projects/bbmap/).

The hybrid read set (both Illumina and ONT reads) for each isolate was assembled using Unicycler 0.4.0 (Wick *et al*. 2017) in its conservative mode. Unicycler performs the assembly of the Illumina reads with SPAdes 3.12.0 (Bankevich *et al*. 2012), and assembly graph scaffolds were then prepared using ONT reads. Unicycler was used to polish the final assembly of Illumina reads, and Pilon (Walker *et al*. 2014) was applied to reduce the rate of small base-level errors. The resulting assembly graph was visualized using Bandage (Wick *et al*. 2015).

The assembled genomes were annotated with the Rapid Annotation using Subsystem Technology (RAST) annotation server (http://rast.theseed.org/FIG/rast.cgi; Aziz *et al*. 2008; Brettin *et al*. 2015; Overbeek *et al*. 2014). Annotation completeness was assessed using BUSCO v3 (Rhizobiales database, Waterhouse *et al*. 2018).

### Comparative genomics

We inferred orthologs from all nine assembled genomes and reference genomes of 15 rhizobial strains (*Mesorhizobium*, *Ensifer*, *Rhizobium*, *Bradyrhizobium*, *Azorhizobium*, *Paraburkholderia* and *Cupriavidus*; Supporting Information Table S3), including both chromosomes and plasmids, using SonicParanoid with the most sensitive parameter settings (Consentino and Iwasaki 2018).

We extracted single-copy orthologs that were found in all nine sequenced strains and two reference strains (*M. japonicus* MAFF303099 and *M. loti* TONO). Orthologous groups were assigned as genes in the core genome or in the symbiosis island based on the genome locations of the reference strain, *M. japonicus* MAFF303099: coordinates of the symbiosis island were 4,643,427 [*intS*] – 5,255,770 [*trnF*-GAA], and the rest was considered to be the core genome (Kaneko *et al*. 2000). We generated a multiple nucleotide sequence alignment of each single-copy orthologous group using MAFFT 7.245 (Katoh and Standley 2013) with the E-INS-i algorithm, and extracted biallelic single nucleotide polymorphisms (SNPs) from the alignments.

To visualize the genetic variations among the nine sequenced strains, we performed principal component analyses (PCA) based on the following datasets: 1) presence/absence of orthologs, 2) the copy number variation of orthologs, 3) SNPs in the core genome and 4) SNPs in the symbiosis island. PCA was performed using the *prcomp* function implemented in R 3.6.1 (R Core Team (2019); http://www.R-project.org/).

To characterize the genome-wide pattern of polymorphisms in these nine sequenced strains, we calculated nucleotide diversity (π values for synonymous and nonsynonymous sites) and Tajima’s *D* for each gene using MEGA-CC (Kumar *et al*. 2012; Tamura *et al*. 2011).

To investigate the phylogenetic relationships of *L. japonicus*-associated symbionts with other nodule bacteria, phylogenetic trees of each single-copy gene were reconstructed using the maximum likelihood method with the program RAxML-NG (Kozlov *et al*. 2019), together with orthologs from reference sequences. Prior to the tree reconstruction, haplotypes were determined for each gene based on nucleotide substitutions and indels. The nucleotide substitution models were selected by the Akaike Information Criterion, as implemented in ModelTest-NG (Darriba *et al*. 2019). The single most likely tree out of 10 search replicates was saved for phylogenetic analyses. The outgroups of each phylogenetic tree were determined using the Graph Splitting method (Matsui and Iwasaki 2019) based on protein sequences of each gene, under the assumption that an outgroup is the most distant operational taxonomic units from the focal group containing *L. japonicus*-associated symbionts. The sensitivity was set to 2.0 in MMseqs2 (Steinegger and Söding 2017), used in the GraphSplitting method.

We then characterized the topologies of maximum likelihood trees of each single-copy gene based on the following criteria: 1) whether *L. japonicus*-associated symbionts (nine sequenced strains in this study, *M. japonicum* MAFF303099 and *M. loti* TONO) formed a single clade; 2) whether *Lotus*-associated symbionts (*L. japonicus*-associated symbionts, *M. loti* NZP2037 and *M. japonicum* R7A) formed a single clade; and 3) whether *Mesorhizobium* strains (nine sequenced strains in this study and other *Mesorhizobium* strains) formed a single clade. This analysis was performed using in-house scripts written in Python 3.

We also performed synteny analysis using the progressiveMauve 2.4.0 program (Darling *et al*. 2010) to identify any structural variations in the symbiosis islands. The symbiosis islands of each genome were first identified by aligning them with that of *M. japonicum* MAFF303099. We then excised the region and aligned the direction using SnapGene software (GSL Biotech; https://www.snapgene.com).

### Cross-inoculation experiments

To quantify the effects of rhizobial symbionts, host plants and their interactions on plant phenotypes, we performed a cross-inoculation experiment using nine sequenced rhizobial strains (Supporting Information Table S1) and 15 *L. japonicus* natural accessions (Supporting Information Table S2), resulting in 135 combinations in total. Seeds of *L. japonicus* accessions were obtained from the Natural BioResource Project.

Prior to inoculation experiments, we prepared inoculant strains in the logarithmic growth phase. We cultured rhizobial strains on a TY agar plate for 4 days at 28°C, and then picked single colonies and precultured them with shaking in 2 mL TY liquid media for 3 days at 28°C. Aliquots (200 μL) of precultured strains were transferred into 50 mL lots of TY liquid medium and cultured with shaking at 28°C for 48 h. The cultured strains were precipitated by centrifugation at 5800 g for 3 min, washed with sterilized water three times and adjusted to 1.0 × 10^7^ cells/mL (based on optical density at 600 nm).

Partly scrubbed *L. japonicus* seeds were surface sterilized by immersion in 2% sodium hypochlorite for 3 min and rinsed three times with sterile distilled water. After overnight imbibition, the swollen seeds were sown onto 0.8% agar plates, incubated in the dark for 3 days at 20°C and then grown at 20°C under 16/8 light/dark conditions for 24 h. The rooting plants were transplanted into Leonard jars (Leonard 1943) filled with 300 mL sterilized vermiculite with 300 mL sterilized nitrogen-free B&D medium (Broughton and Dilworth 1971) and grown at 20°C under the same lighting conditions for 3 days. Finally, we inoculated 20 mL of each concentration-adjusted rhizobial strain into Leonard jars and grew them at 20°C under the same lighting conditions for 21 days. We then harvested whole plant bodies, imaged all individuals with a high resolution scanner and separated them into shoots and roots. The shoots were dried over 48 h at 65°C, and then the dry weights were measured. For root phenotypes, we measured the numbers and areas of nodules from the scanned data. Shoot dry weight (SDW in g), number of nodules (NON), total size of nodule (NOA in mm²) and nodule size per nodule (NOA/NON ratio defined as NOAi in mm²) were obtained from all individuals used in the experiment. We repeated all inoculation experiments twice. When we grew *japonicus* without inoculation, plants did not form any nodules and lost most of the leaves (data not shown).

### Data analysis of the inoculation experiments

We performed analysis of variance to test whether genotypes of inoculant symbiont, those of host plants and their interactions (G × G) significantly influenced the four measured phenotypes (SDW, NON, NOA and NOAi). We considered mean values of each rhizobial strain (means of phenotypes of 15 plant accessions) as the rhizobial quality. To quantify G × G interactions, we generated Euclidean distance matrices based on the phenotypic differences between rhizobial strains by using the standardized phenotypic values whose mean values of each rhizobial symbiont were set to 0, thereby controlling for bacterial genetic effects. We used mean phenotype values and Euclidean distance matrices for the following association analyses with the bacterial genome sequences. R v. 3.6.1 was used for analysis of variance (R Core Team 2019), and the Euclidean distance was calculated by using SciPy (Virtanen *et al*. 2019).

To determine the genomic regions associated with rhizobial quality, we first performed an SNP-based association analysis that examined the correlation between genome-wide SNPs and rhizobial quality using a linear model implemented by LIMIX (Lipert *et al*. 2014). We then performed Mantel tests to evaluate whether the genetic distance between rhizobial strains was correlated positively with the difference in the variations of rhizobial quality or those of G × G interactions. We performed these analyses for the core genome and symbiosis islands separately to examine which genomic regions were more strongly associated with variations in rhizobial quality. For the genetic distance, kinship matrices were calculated using mixmogam (Segura *et al*. 2012). Mantel tests were conducted using Spearman’s rank correlation implemented in the scikit-bio v. 0.5.4 (http://scikit-bio.org/) library of Python.

## RESULTS

### Bacterial genome sequencing

Nine rhizobial genome sequences were obtained using ONT MinION and Illumina HiSeq sequencing. From MinION sequencing, 1,011,067 reads (mean length 6748 bp) were obtained, covering a total of 6.823 Gb. After pre-processing the MinION sequences, 822,488 reads were allocated into the nine samples, which ranged from 47,958 to 142,669 in read number and 370–1181 Mb in total length (Supporting Information Table S4). From HiSeq sequencing, 32,511,368 pre-processing reads were obtained and allocated into the nine samples, which ranged from 2,432,754 to 4,476,543 in read number and 364–671 Mb in total length. All quality-filtered reads from both the MinION and HiSeq sequences were used for the *de novo* assembly analyses.

### *De novo* assembly and annotation

Nearly complete assembled genomes of all nine rhizobial strains were obtained using Unicycler (Wick *et al*. 2017) by combining MinION long reads and HiSeq short reads. All nine rhizobial strains have large circular genomes, considered as chromosomes (6.652–8.451 Mb; Fig. 1 and Supporting Information Table S1). Five of the nine strains had several shorter plasmid-like circular genomes (56–654 kb), and two of the strains had short fragment sequences, which were presumably contaminants (< 10 kb fragment lengths and not closely related to *Mesorhizobium*). The total genome sizes excluding presumable contaminant fragments ranged from 7.108 to 8.451 Mb. The genome sizes of all assembled genomes were similar to that of the reference strain, *M. japonicum* MAFF303099 (chromosome, 7.036 Mb; pMLa, 0.351 Mb, pMLb, 0.251 Mb; and total 7.596 Mb). All genomes without presumed contaminant fragments were used for the following analyses. Gene predictions from all assembled genomes were conducted using the RAST server (Overbeek *et al*. 2014). The number of coding sequences was 7173–8377, including six rRNA genes and 50–61 tRNA genes in each genome (Supporting Information Table S1). The number of coding sequences was also similar to that of *M. loti* MAFF303099 (7343 genes). The evaluation of completeness of gene prediction by BUSCO analyses showed that all nine rhizobial strains had no missing BUSCO. A few were fragmented (0.292–1.17% fragmented) and most BUSCOs were complete (98.8–99.7% complete; Supporting Information Table S1). These BUSCO analyses indicated that we had successfully reconstructed nearly complete genomes of nine rhizobial strains.

**Figure 1.**
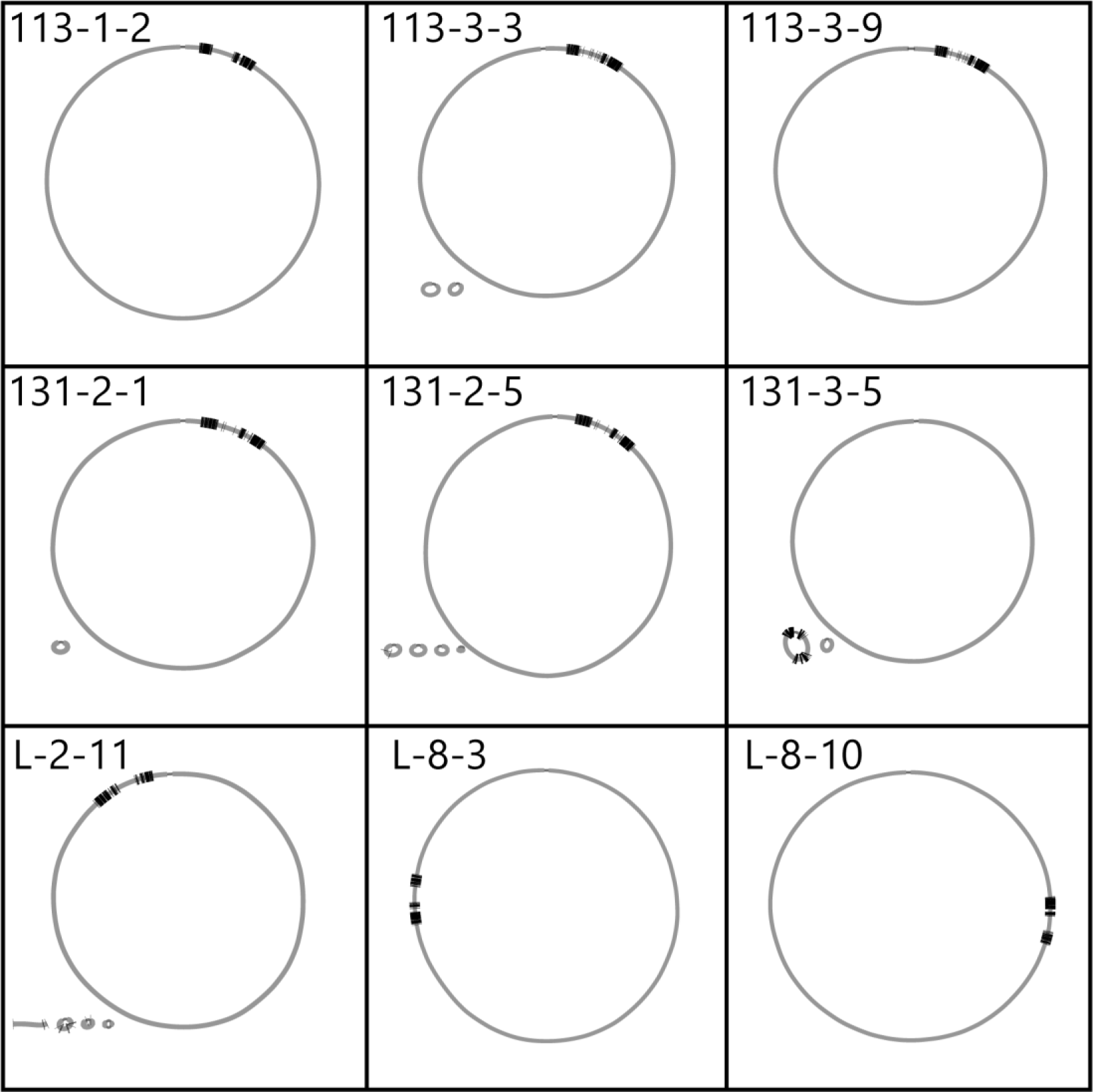
Assembly graphs of nine newly assembled rhizobial genomes. Gray circles and bars indicate assembled genomes. Thick black bars on the genomes indicate genes identified as homologs of genes on the symbiosis island of the reference strain, *M. japonicum* MAFF303099.

### Ortholog analysis

The newly assembled genomes, including both chromosomes and plasmids, were compared with the reference genomes of 15 rhizobial strains (Supporting Information Table S3). We identified a total of 15,712 orthologous groups (OGs) of genes using SonicParanoid (Consentino and Iwasaki 2018). Among these OGs, 3095 were conserved among 11 *L. japonicus*-associated symbionts (*M. japonicum* MAFF303099, *loti* TONO and nine newly assembled genomes), and 3047 OGs were conserved among 13 *Lotus*-associated symbionts (the same 11 *L. japonicus*-associated symbionts and two *L. corniculatus*-associated symbionts, *M. japonicum* R7A and *M. loti* NZP2037).

We obtained 2239 OGs as single-copy orthologs found in all of nine sequenced genomes of symbionts and two reference strains (*M. japonicum* MAFF303099 and *M. loti* TONO; Supporting Information Table S5). Seventy-six single-copy orthologs were located in symbiosis islands based on the coordinates of *M. japonicum* MAFF303099.

Using all the identified OGs, we investigated the presence/absence of genes present on the symbiosis islands of *M. japonicum* MAFF303099 and reported to be related to symbiotic features such as nitrogen fixation, nodulation factor assembly and protein secretion systems (Porter *et al*. 2019; Souza *et al*. 2012; Wang *et al*. 2014). All 28 nitrogen fixation-related genes (*nif*, *fdx* and *fix*) were present in the nine newly assembled genomes and in other genomes of *Lotus*-associated symbionts (Fig. 2).

**Figure 2.**
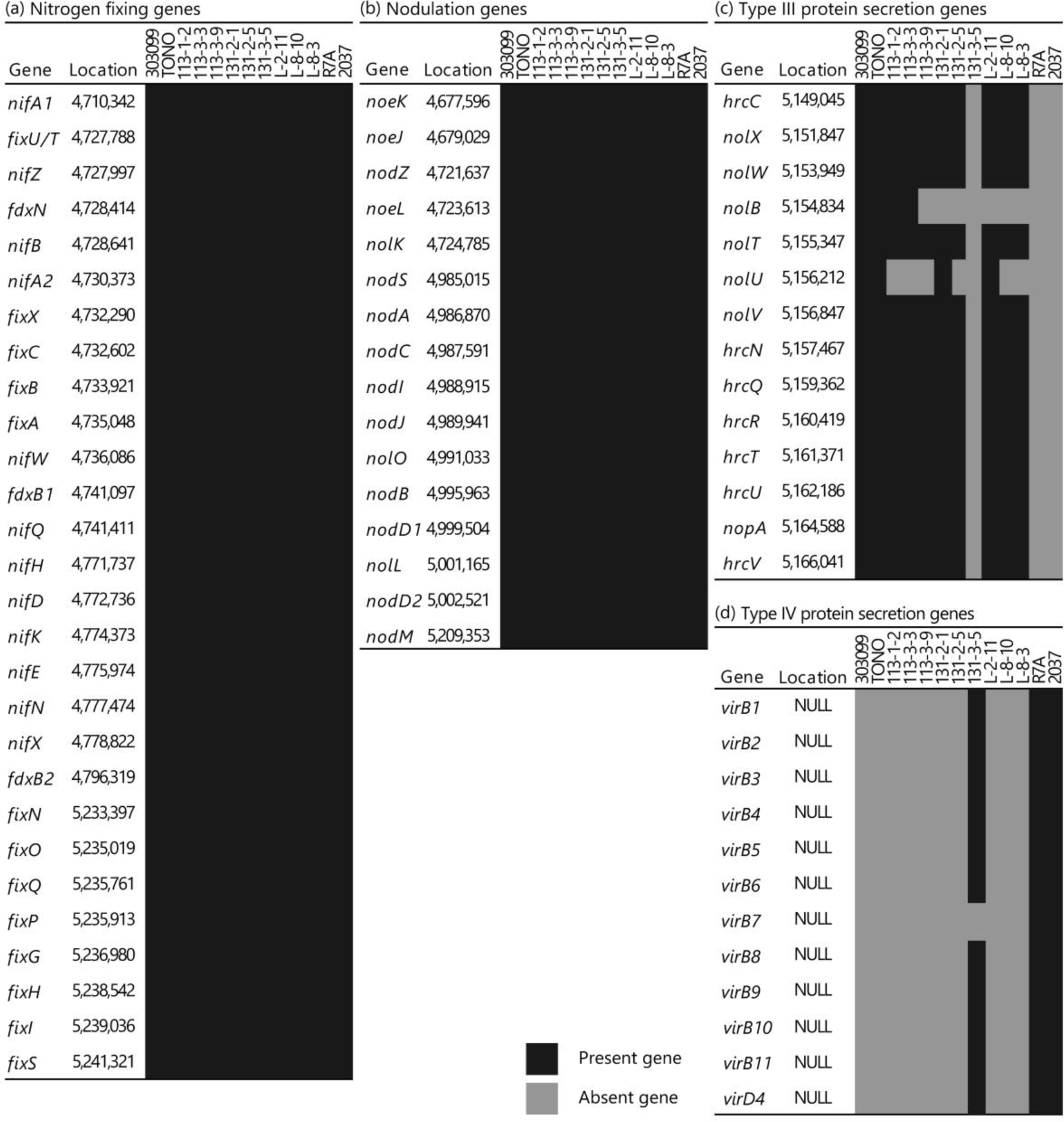
Presence/absence of variations among genes reported to be related to the symbiosis: (a) nitrogen-fixing genes; (b) nodulation genes; (c) type III protein secretion system genes; and (d) type IV protein secretion system genes. Each row indicates rhizobial strains with abbreviations as follows: 303099, *M. japonicum* MAFF303099; TONO, *M. loti* TONO; R7A, *M. japonicum* R7A; and 2037, *M. loti* NZP2037. Black and gray cells indicate the presence or absence of genes, respectively.

Sixteen nodulation factor assembly genes (*nod*, *nol* and *noe* genes) were present in all nine genomes. All 14 type III protein secretion system (T3SS) genes were absent in one *L. japonicus*-associated strain (131-3-5) and in two *L. corniculatus*-associated strains; two of them (*nolB* and *nolU*) were also missing in multiple genomes of *Lotus*-associated symbionts. By contrast, one *L. japonicus*-associated strain (131-3-5) had nearly the complete set of the type IV protein secretion system (T4SS) genes that were absent in other *L. japonicus*-associated symbionts, except for the *virB7* gene.

### Principal component analysis (PCA)

PCA showed genome-wide genetic variations between strains. When based on the presence/absence of orthologs, PC1 and PC2 explained 49.41% and 14.46%, respectively (Supporting Information Fig. S1A), When based on copy number variations, PC1 and PC2 explained 38.05% and 25.16%, respectively (Supporting Information Fig. S1B). In both plots, there were two closely related pairs of strains (131-2-5 and 131-3-5, L-8-3 and L-8-10) and no clear geographical clusters.

We extracted biallelic SNPs from each gene group (core genome 528,912 bp, symbiosis island: 2497 bp) and then performed PCA separately (Fig. 3A, B). PCA based on the SNPs in the core genome showed similar patterns to the one generated by genome-wide ortholog profiles (Fig. 3A); there were two closely related pairs of strains, which were also observed in the ortholog-based plot. By contrast, PCA using the SNPs of the symbiosis island showed a distinct pattern (Fig. 3B): we again found two pairs of closely related strains, but they were different from those observed in the core genome or genome-wide ortholog profiles (113-3-3 and 113-3-9, 131-2-1 and 131-2-5). Overall, PCA suggested that the pattern of relatedness between strains differed markedly between the core genome and the symbiosis island.

**Figure 3.**
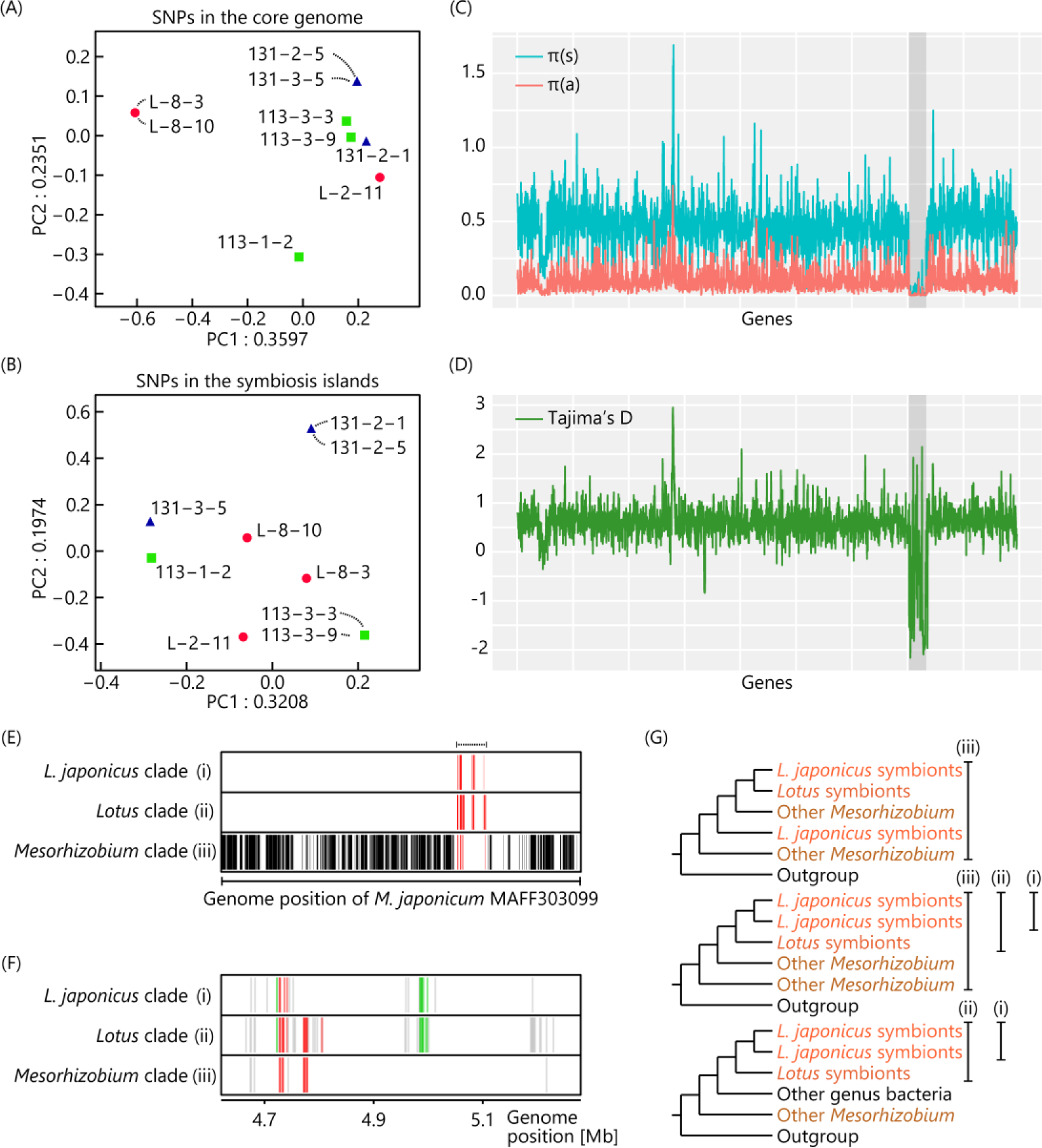
Genomic outlines of nine sequenced strains. (A, B) Principal component analyses of nine sequenced strains using single nucleotide polymorphisms (SNPs) in the core genome (A) and in the symbiosis island (B). Forms and colors of dots indicate sampling localities of each rhizobial symbiont with the strain type designated in brackets: blue triangle, Aomori (131); green square, Tottori (113); and red circle, Miyakojima (L). (C, D) Genome-wide distribution of nucleotide diversity statistics π (C) and Tajima’s *D* (D), calculated for each single-copy gene described in Supporting Information Table S4. The order of genes was based on the genome assembly of *M. japonicum* MAFF303099. The shaded region indicates the symbiosis island. (C) Red and blue lines indicate the nucleotide diversity statistic π at nonsynonymous and synonymous sites, respectively. (E, F and G) Comparison of phylogenetic tree topologies in the rhizobial whole genomes (E) and the symbiosis island (F). Bars are indicated if: (i) *L. japonicus*-associated symbionts (nine sequenced strains, *M. japonicum* MAFF303099 and *M. loti* TONO) form a single clade; (ii) *Lotus*-associated symbionts (*L. japonicus*-associated symbionts and *M. loti* NZP2037, and *M. japonicum* R7A) form a single clade; and (iii) *Mesorhizobium* strains (nine sequenced strains in this study and other *Mesorhizobium* strains) form a single clade. (E) The black and red bars indicate genes on the core genome and the symbiosis island, respectively. (F) Each red, green and gray bar indicates nitrogen-fixing, nodulation and unknown genes. (G) Schematic trees showing clades satisfying criteria i–iii listed above.

### Genome-wide view of polymorphisms and gene genealogies

To understand the genome-wide landscape of polymorphisms, we calculated nucleotide diversity and Tajima’s *D* statistic for each gene across the genome (Fig. 3C, D; Supporting Information Table S5). Genes in the core genome showed markedly higher nucleotide diversity at both synonymous and nonsynonymous sites (mean 0.4953 and 0.1029, respectively) compared with those of genes on the symbiosis island (mean 0.0250 and 0.006367, respectively). This pattern is consistent with an observation of the smaller number of genes in a previous study (Bamba *et al*. 2019b), and indicates a signature of horizontal gene transfer (HGT) of the symbiosis island. A steep change in diversity at the borders of the symbiosis island suggests that it would have behaved as a unit of HGT.

Tajima’s *D* statistic of the core genome (mean 0.6990) was also markedly higher than that of the symbiosis island (mean –0.5304; Fig. 3D; Supporting Information Table S5). It is also important to note that Tajima’s *D* statistic of the core genome was generally positive, indicating the excess of alleles with intermediate frequencies. This pattern would be expected if there were a clear population structure in the core genome, which was indeed observed in our PCA (Fig. 3A). By contrast, Tajima’s *D* statistic of the symbiosis island was generally negative, indicating an excess of rare alleles in this region, and to be expected under a scenario of recent selective sweeps including HGT.

Patterns of phylogenetic relationships were also distinct between the core genome and the symbiosis island (Fig. 3E–G, Supporting Information Table S5). In the core genome, the majority of the gene trees (1555/2163 genes) showed a single *Mesorhizobium* clade, whereas there was no gene tree in which *L. japonicus-* or *Lotus*-associated symbionts formed a clade (Fig. 3E; Supporting Information Table S5). By contrast, in the symbiosis island, 14/76 gene trees showed the *L. japonicus*-associated symbionts clade, 55 genes formed the *Lotus*-associated symbionts clade and 14 genes showed the *Mesorhizobium* clade (Fig. 3F; Supporting Information Table S5). These results suggest that *L. japonicus*-associated symbionts mostly have the core genome of *Mesorhizobium*, but their symbiosis island clearly has a different evolutionary origin, supporting the HGT of the symbiosis island into the diverse genetic background of *Mesorhizobium*. In addition, multiple types of topologies in the symbiosis island indicated a history of recombination within the symbiosis island.

### Structural variations in the symbiosis island

An alignment by progressiveMauve analysis (Darling *et al*. 2010) provided the genomic regions corresponding to the symbiosis island of the reference strain, *M. japonicum* MAFF303099. All of the nine newly assembled genomes harbored the symbiosis island. Eight had the symbiosis island on the chromosome, but in strain 131-3-5, the symbiosis island was identified on the plasmid (Fig. 1).

Alignment by progressiveMauve showed that the synteny of symbiosis island of *L. japonicus*-associated symbionts was mostly conserved. Five conservative synteny blocks were identified in the symbiosis island, and all blocks were found in eight of the nine sequenced strains, except for strain 131-3-5, which lacked the whole type III protein secretion system gene cluster (*nol–hrc*). This was consistent with an ortholog search using SonicParanoid (Consentino and Iwasaki 2018), which revealed the lack of all *nol*–*hrc* genes (Fig. 2).

The largest synteny block, 21 kb away from the start positions of symbiosis islands (symbiosis islands integrase; *intS*) on *M. japonicum* MAFF303099 (Fig. 4A), contained nitrogen-fixing genes (*nif*, *fix* and *fdx*) and several nodulation genes (*nod*, *noe* and *nol*). We note that this largest block was inverted in the strain L-2-11. Between the *nif–fdx* and *nod–nol* blocks, there was a hypervariable region containing many transposase insertions (around 135 kb in *M. japonicum* MAFF303099).

**Figure 4.**
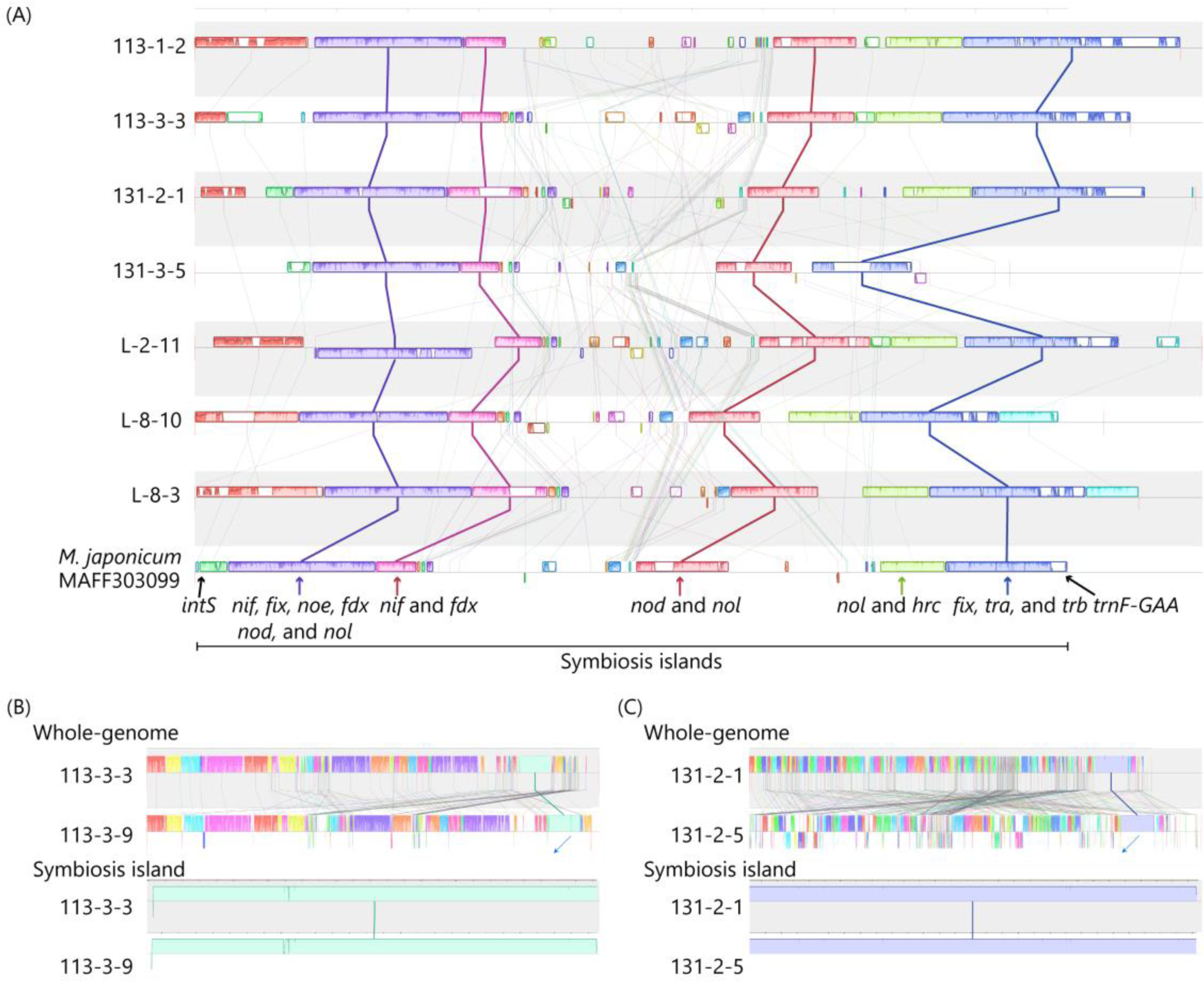
An alignment of the symbiosis islands of *L. japonicus*-associated symbionts, using progressiveMauve software. Bordered and connected boxes indicate similar sequence compositions among sequences. (A) The boxes connected by bold lines indicate conservative genetic clusters among all *L. japonicus*-associated symbionts. The purple and pink/purple conserved blocks, red conserved block and blue and yellow conserved blocks are referred as *nif–fdx* blocks, *nod* blocks and *nol–hrc* blocks in the main text, respectively. (B, C) Alignment of two pairs of rhizobial strains harboring almost identical symbiosis islands, 113-3-3 and 113-3-9 (B) and 131-2-1 and 131-2-5 (C), generated by progressiveMauve software. The upper and lower diagrams show alignments of whole genomes and the symbiosis islands, respectively. Boxes connected by bold lines indicate the symbiosis islands.

We found that two pairs of strains harbored extremely similar symbiosis islands, even including the hypervariable region (113-3-3 and 113-3-9, 131-2-1 and 131-2-5; Fig. 4B, C). The sequence identity values between 113-3-3 and 113-3-9 and between 131-2-1 and 131-2-5 were 99.6% and 99.9%, respectively. These two pairs corresponded to the pairs identified in the PCA of SNPs from the symbiosis island (Fig. 3B). Because their core genomes are highly different, these data also provide strong evidence for recent HGT in the whole symbiosis island.

### Cross-inoculation experiments

We performed a cross-inoculation experiment using 15 *L. japonicus* accessions and nine rhizobial symbionts, resulting in a total of 135 combinations (Supporting Information Tables S1 and S2). We obtained four phenotypes (SDW, NON, NOA and NOAi) from 1189 individuals (5–14 per combination; Fig. 5). All phenotypic traits were significantly correlated with each other (Supporting Information Figs S2 and S3, all Pearson’s product-moment correlation *P* < 2e^−16^). All correlation coefficients were positive (0.246 to 0.637), except for that between NON and NOAi (–0.351).

**Figure 5.**
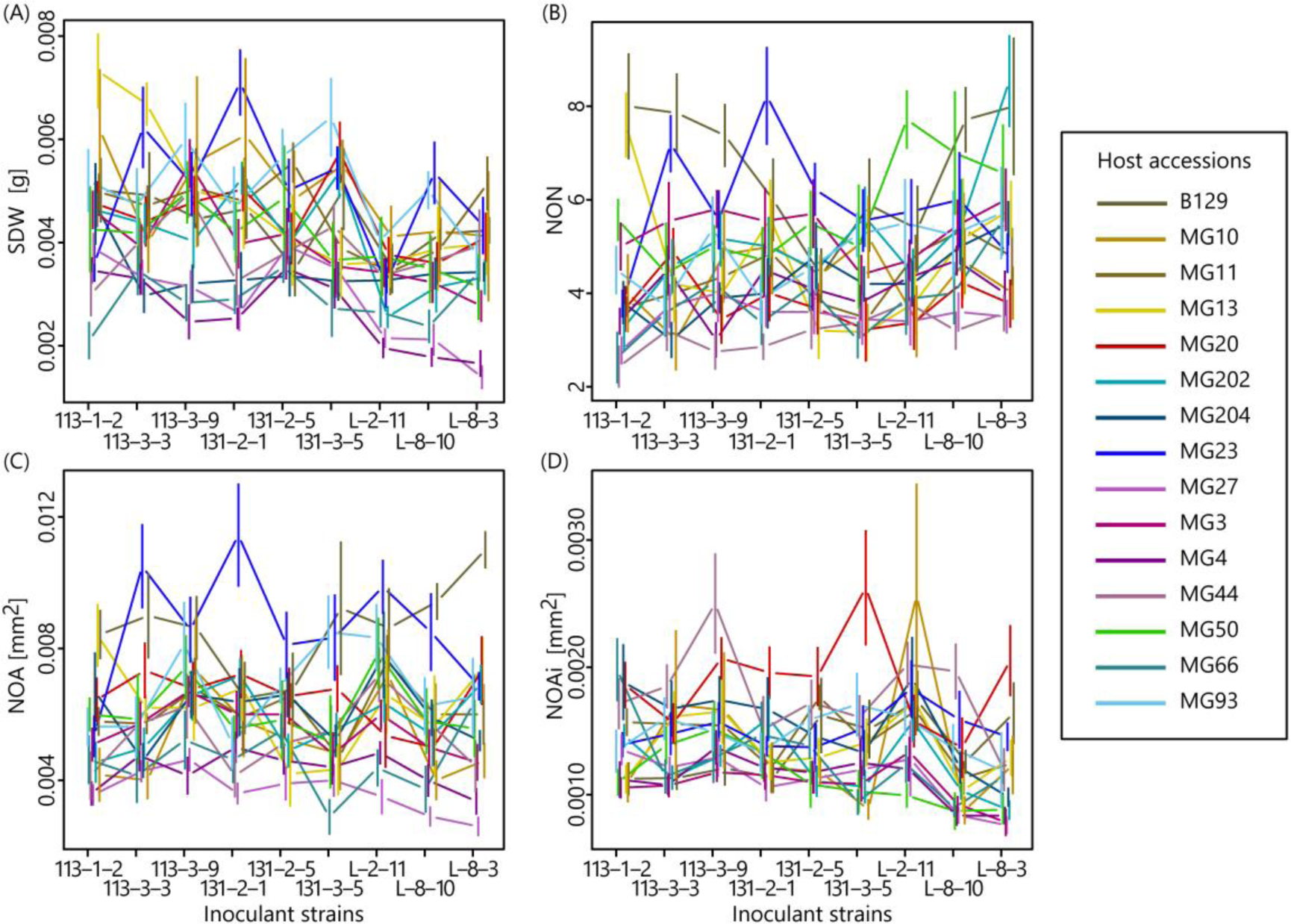
Plant phenotypes of the cross-inoculation experiments, (A) shoot dry weight (SDW), (B) nodule numbers (NON), (C) total nodule size (NOA), (D) nodule size per nodule (NOAi). Mean values and standard errors (bars) are shown.

In the cross-inoculation experiment, we detected significant effects of host, symbionts and their interactions on all four phenotypes (Table 1: all host, symbiont and interaction effects were significant at *P* < 0.001). For all phenotypes, the effect of host was the largest (partial η^2^ = 0.200–0.268), followed by the host × symbiont interaction effect (partial η^2^ = 0.145–0.168) and the effect of symbionts (partial η^2^ = 0.038–0.081; Table 1), indicating that the host phenotypes were more strongly affected by host × symbiont interactions than the sole symbiont effect. Furthermore, it is worth noting that the combinations of hosts and symbionts from the same localities did not necessarily show higher phenotypic values than nonnative combinations (Supporting Information Table S6), which was not consistent with the pattern expected from local adaptation.

To understand what genes or genomic regions of rhizobial symbionts could be responsible for variations in partner quality, we first quantified the rhizobial quality (Supporting Information Fig. S2) and G × G effects (Supporting Information Fig. S4). There were 1.233–1.413 times greater differences between minimum and maximum values of variations in partner quality.

Genes previously reported as being involved in symbiosis would be obvious candidates explaining the rhizobial variations in partner quality. Therefore, we first investigated the correlations between the variations in partner quality (represented by mean phenotypic values) and the presence/absence variations of these symbiosis genes. Analysis of variance using the genes in which we found the presence/absence variations (Fig. 2) revealed that none were significantly correlated with variations in partner quality (*P* > 0.05; Supporting Information Table S7). Next, we investigated the correlation between the variations in partner quality and genetic distances of genes that were reported as candidate genes responsible for rhizobial quality (Klinger *et al*. 2016). Mantel testing using these candidate genes found that none were significantly correlated with rhizobial quality variations (Supporting Information Table S8). These results suggest that known symbiosis-related genes would not be responsible for the rhizobial variations in partner quality detected in the cross-inoculation experiment.

We then performed an SNP-based association analysis between rhizobial genomes and rhizobial variations in partner quality. We found that SNPs in the core genome were more strongly associated with rhizobial quality than those in the symbiosis islands (Supporting Information Fig. S5). In the linear model for SDW, NON and NOAi, *P* value distributions were strongly skewed toward small values in the core genome (62.6%, 70.7% and 58.5% of SNP *P* values were < 0.05, respectively; Supporting Information Fig. S5A–D). These skewed *P*-value distributions in the core genome presumably resulted from population structure and strong linkage disequilibrium. On the other hand, *P* value distributions were not strongly skewed in the symbiosis islands. Thus, the *P*-values were < 0.05 in 10.8% of SDW SNPs, 5.3% of NON SNPs, 11.8% of NOA SNPs and 8.8% of NOAi SNPs; Supporting Information Fig. S5E–H).

Next, we performed Mantel tests to determine whether the genetic distance between rhizobial strains was positively correlated with variations in partner quality. The Mantel test results between the variations of rhizobial quality for SDW and NOAi were significantly correlated with the reciprocal of rhizobial core genome kinships (*P* < 0.05 after Bonferroni corrections; Table 2). By contrast, we did not recognize any correlation between variations in rhizobial quality and the kinships of symbiosis islands. These results suggest that variation in the core genome, rather than in the symbiosis islands, could explain the rhizobial variations in partner quality.

Variations in G × G interactions were not significantly correlated with rhizobial core genomes or symbiosis islands (Supporting Information Table S9). However, it is noteworthy that there were variations in G × G interactions between the pairs that appeared to have almost identical symbiosis islands but highly different core genomes (Fig. 4B, C; 113-3-3 and 113-3-9: 131-2-1 and 131-2-5; Supporting Information Fig. S4). There were no significant correlations for G × G variations between these symbiont pairs, except for NOAi of 113-3-3 and 113-3-9 (*P* < 0.05 after Bonferroni corrections; Supporting Information Table S10). Because the symbiosis islands are nearly identical, the observed G × G variations should due to genetic variation in the core genome.

## DISCUSSION

### *De novo* assembled genomes of *Lotus japonicus*-associated nodule bacteria

While there have been several attempts to sequence the whole genomes of nodule bacteria (Kaneko *et al*. 2000, 2002; Amadou *et al*. 2008; Lee *et al*. 2008; Reeve *et al*. 2010ab, 2015; Ramsay *et al*. 2013; Moulin *et al*. 2014; Wang *et al*. 2014; Shimoda *et al*. 2016; Nagymihály *et al*. 2017; Liang *et al*. 2018), *de novo* sequencing of multiple strains associated with a single plant species has been uncommon. Here, we performed a whole-genome sequencing analysis of nine *L. japonicus-*associated symbionts by exploiting both long-read (Oxford Nanopore MinION) and short-read (Illumina HiSeq) sequencers, which enabled us to generate high quality *de novo* assembled genomes for each strain.

Comparative genomic analyses of these sequenced genomes provided clear evidence for HGT of the symbiosis island. First, patterns of phylogenetic relationships were distinct between the core genome and the symbiosis island (Fig. 3E, F; Supporting Information Table S5). In the core genome, there was no gene forming an *L. japonicus-* or *Lotus*-associated symbiont clade, but in the symbiosis island, 14 and 55 of 76 genes showed *L. japonicus*- or *Lotus*-associated symbiont clades, respectively, indicating that the evolutionary origin of the symbiosis island was clearly different from that of the core genome (Fig. 3E, Supporting Information Table S5). Second, we observed a marked decline in the nucleotide diversity π statistic at synonymous and nonsynonymous sites and negative Tajima’s *D* statistic, which are also evidence for HGT (Fig. 3C, D). A steep change in diversity at the borders of the symbiosis island suggests that it would have behaved as a unit of HGT, while the multiple topologies of gene trees in the symbiosis island also indicate an evolutionary history of internal recombination events. Third, there was a pair of strains harboring almost identical sequences of the symbiosis island, but with highly distinct core genome backgrounds, which is also a signature of recent HGT (Fig. 4B, C). Including our previous study of *L. japonicus*-associated nodule bacteria (Bamba *et al*. 2019b), there have been several studies demonstrating HGT of symbiosis islands based on sequences of a few genes (Barcellos *et al*. 2007; Steenkamp *et al*. 2008; Menna and Hungria 2011; Koppell and Parker 2012; Parker and Rousteau 2014; Lemaire *et al*. 2015; Bamba *et al*. 2016). Thus, our *de novo* assembled genome data have provided multiple signatures of HGT at an unprecedented level of resolution.

The assembled genomes also revealed blocks of conserved synteny as well as extensive structural rearrangements in the symbiosis island. While Shimoda *et al*. (2016) showed that there were three conserved regions (*nif*, *nod* and type III protein secretion system) in the symbiosis islands of *Lotus*-associated symbionts, we found five conserved blocks and many rearrangements, including an inversion and loss of genes and gene clusters. As these structural rearrangements were found in the symbiosis island of bacteria associated with a single legume species, rearrangements or gene gain/loss could occur over a short time. We also found evidence of recombination within the symbiosis island based on the topologies of gene trees (Fig. 3E, F). Signatures of gene gain/loss and recombination have also been reported for several rhizobial genera (*Burkholderia*, De Meyer *et al*. 2016; *Bradyrhizobium*, Sugawara *et al*. 2013; Bouznif *et al*. 2019; Porter *et al*. 2019), and the genomic insight of the symbiosis island of *L. japonicus*-associated symbionts is consistent with the emerging perspective that recombination and gene gain/loss are not rare events in rhizobial symbiosis islands/plasmids (Porter *et al*. 2019).

### Genomic regions associated with variations in partner quality

By integrating the data of cross-inoculation experiments and assembled whole genomes, we investigated which genomic regions could be responsible for variations in partner quality. We found that plant growth was significantly influenced by host genotypes, symbiont genotypes and host × symbiont (G × G) interactions (Table 1). We then examined whether rhizobial variations in partner quality were explained by the following genetic factors: (i) presence/absence variation of symbiosis genes, (ii) SNPs in the rhizobial genomes and (iii) genetic kinships of the core genome and the symbiosis island. We found that the rhizobial core genome variations explained the rhizobial variations in partner quality: Mantel tests showed that the quality variations in rhizobial symbionts and their core genome kinship were significantly correlated in two phenotypes (SDW and NOAi; Spearman rank correlations *P* < 0.05 after Bonferroni correction; Table 2), whereas the kinship of the symbiosis island was not. In the SNP-based association analyses, *P*-value distributions of core genomes SNPs were strongly skewed toward small values (Supporting Information Fig. S5), suggesting that many SNPs across the genomes are correlated with variations in partner quality, consistent with the results of Mantel tests. Such strongly skewed *P*-value distributions possibly arise from extensive genome-wide linkage disequilibrium. Bacterial genomes generally show strong linkage disequilibrium given their asexual reproduction (Chen *et al*. 2015), which could make it difficult to pinpoint the responsible regions/genes on the genome using association-based analysis.

Our finding of significant correlations between core genome variations and the rhizobial variations in partner quality might partly explain why such variations persist in legume–rhizobia mutualisms. In *L. japonicus*-associated symbionts, massive genetic variations in the core genome would be maintained by recurrent HGT of the symbiosis islands into diverse *Mesorhizobium* bacterial strains (Bamba *et al*. 2019b). Local *Mesorhizobium* communities could thus serve as a source of standing genetic variation of core genomes, which might prevent variations in partner quality from fixing even under the presence of selection favoring high-quality partners: i.e. a stabilizing mechanism in mutualisms (Heath and Stinchcombe 2014). In the context of the mutation–selection balance model (Van Dyken *et al*. 2011; Smith *et al*. 2014), our study serves to illuminate the role of HGT among symbiosis islands that indirectly supply mutations in core genomes contributing to variations in partner quality.

A few studies have reported that variations in the symbiosis islands explain rhizobial variations in partner quality, unlike our finding in *L. japonicus*-associated bacteria (Klinger *et al*. 2016; Porter *et al*. 2019). Klinger *et al*. (2016) showed that variations in partner quality are explained by variations in the *nifH*, *nifA*, *fixC*, *nodB* and Rleg_4928 (*fixB*) genes. Although all these genes are present in the genomes of *L. japonicus-*associated symbionts (Fig. 2), genetic distances of these genes among strains and variations in rhizobial quality were not significantly correlated, suggesting that these genes do not explain the variations in partner quality in our system (Supporting Information Table S8). Porter *et al*. (2019) showed that the absence of rhizobial symbiosis genes lessens the quality of rhizobial symbionts. However, in our nine sequenced rhizobial genomes, none of the symbiotic genes showing the presence/absence of variations—all type III protein secretion system-related genes (Fig. 2)—were significantly correlated with rhizobial quality variations (*P* < 0.05 after Bonferroni correction; Supporting Information Table S7). We note that type III protein secretion system-related genes might not be involved in interactions between *L. japonicus* and *Meshorhizobium* because dysfunctional mutants of such genes in *M. japonicum* did not show phenotypic changes in terms of nodule-forming ability for *L. japonicus* B129 (Okazaki *et al*. 2010), although their effects have been detected in other *Lotus* species (Okazaki *et al*. 2010; Mercante *et al*. 2015).

We speculate that the contrasting findings between our study and previous ones might have arisen in part from differences in sampling schemes and in the unique history of *L. japonicus*-associated symbionts. Both Klinger *et al*. (2016) and Porter *et al*. (2019) focused on rhizobial populations harboring similar core genomes. Porter *et al*. (2019) used 38 strains possessing a similar core genome as a recombining population, and Klinger *et al*. (2016) analyzed strains collected from nitrogen-enriched experimental fields, and the nucleotide diversity of their core genomes was as low as that of the symbiosis islands. By contrast, data compilation by Bamba *et al*. (2019b) revealed that one of the notable characteristics of the *L. japonicus*-associated rhizobia in Japan is the presence of highly diverse core genomes and the extremely low nucleotide diversity of the symbiosis island. This possibly reflects recent and recurrent HGT of the symbiosis island associated with the population expansion of *L. japonicus* into the Japan archipelago over several thousand years (Bamba *et al*. 2019b). Therefore, it is possible that the symbiosis islands of *L. japonicus*-associated bacteria analyzed in this study have experienced genetic bottlenecks when *L. japonicus* migrated into the Japan archipelago and are too homogeneous to serve as a source for variations in partner quality.

### G × G interactions and local adaptation

In our cross-inoculation experiment, we found that plant growth was significantly influenced by host × symbiont (G × G) interactions, even more than symbiont genotypes (Table 1), as is also observed in a few other symbiosis systems (Heath and Tiffin 2007; Barrett *et al*. 2016). Such G × G interactions are suggested to underlie the selective explanations for the persistence of variations in partner quality in mutualisms (Heath and Stinchcombe, 2014), and our cross-inoculation experimental results support this scenario.

We found complex G × G interactions even between two rhizobial pairs sharing almost identical symbiosis islands (113-3-3 vs. 113-3-9; 131-2-1 vs. 131-2-5; Fig. 4B, C and Supporting Information Fig. S4), strongly suggesting that such variations have arisen in part from variations in rhizobial core genomes. Previous studies on G × G interactions in legume–rhizobia mutualisms used relatively few genes, so it remained unclear which genes/genomic regions were responsible for such interactions (Heath and Tiffin 2007; Barret *et al*. 2015;). Here, we provide clear evidence supporting the contribution of rhizobial core genome variations. However, we did not observe statistically significant correlations between the variations in G × G interactions and rhizobial genomic variations based on the Mantel test (Supporting Information Table S9). While G × G interactions should have a genetic basis (Table 1), they might be governed by polygenic factors that would not be detectable at our relatively small experimental scale (15 rhizobia × 9 legume combinations).

Such G × G interactions in a spatial context have been a hotly debated issue in legume–rhizobia mutualisms (Heath and Tiffin 2007; Heath 2010; Porter *et al*. 2011; Heath *et al*. 2012; Ehinger *et al*. 2014; Harrison *et al*. 2017). The geographic mosaic theory of coevolution states that the outcome of reciprocal selection between a particular genotype of one species and a genotype of an interacting species will differ among ecologically distinct locations (Thompson 1994, 1997, 2005; Forde *et al*. 2004; Decaestecker *et al*. 2007; Laine *et al*. 2014;). According to this theory, if legume– rhizobia mutualistic interactions coevolve locally, native rhizobial genotypes are expected to outperform foreign rhizobial genotypes when associated with host genotypes originating from the same locations. In our experiments, the NOA and SDW data can be considered fitness proxies (Ratcliff *et al*. 2012; Younginger *et al*. 2017), but the combinations of hosts and symbionts from the same localities did not show higher phenotypic values in either of them (Supporting Information Table S6), which is not consistent with the pattern expected from local adaptation. As is also discussed for the maintenance of variations in partner quality, recurrent HGT of the symbiosis islands into diverse *Mesorhizobium* core genomes might explain in part the absence of local adaptations in the *L. japonicus* associated-symbionts, given that core genome variations are strongly correlated with rhizobial variations in partner quality. With local and recurrent HGT of symbiosis islands, core genome variations underlying variations in partner quality might be prevented from fixation even under the presence of local reciprocal selection between plants and rhizobia. By integrating the full-genome sequencing of rhizobial strains and cross-inoculation experiments, this study has demonstrated a scenario of how variations in partner quality could be maintained in the presence of selection and HGT of symbiosis islands. More genetic studies from a plant perspective would be valuable for the understanding of coevolution between plants and rhizobia. This has now become possible for *L. japonicus*, where full-genome resequenced data have become available for hundreds of natural accessions (Shah *et al*. 2020).

## ACKNOWLEDGMENTS

Wild accessions of *L. japonicus* used in this research were provided by the National BioResource Project “Lotus/Glycine” of the Ministry of Education, Culture, Sports, Science and Technology, Japan.

## SUPPLEMENTARY FIGURES

**Figure S1.**
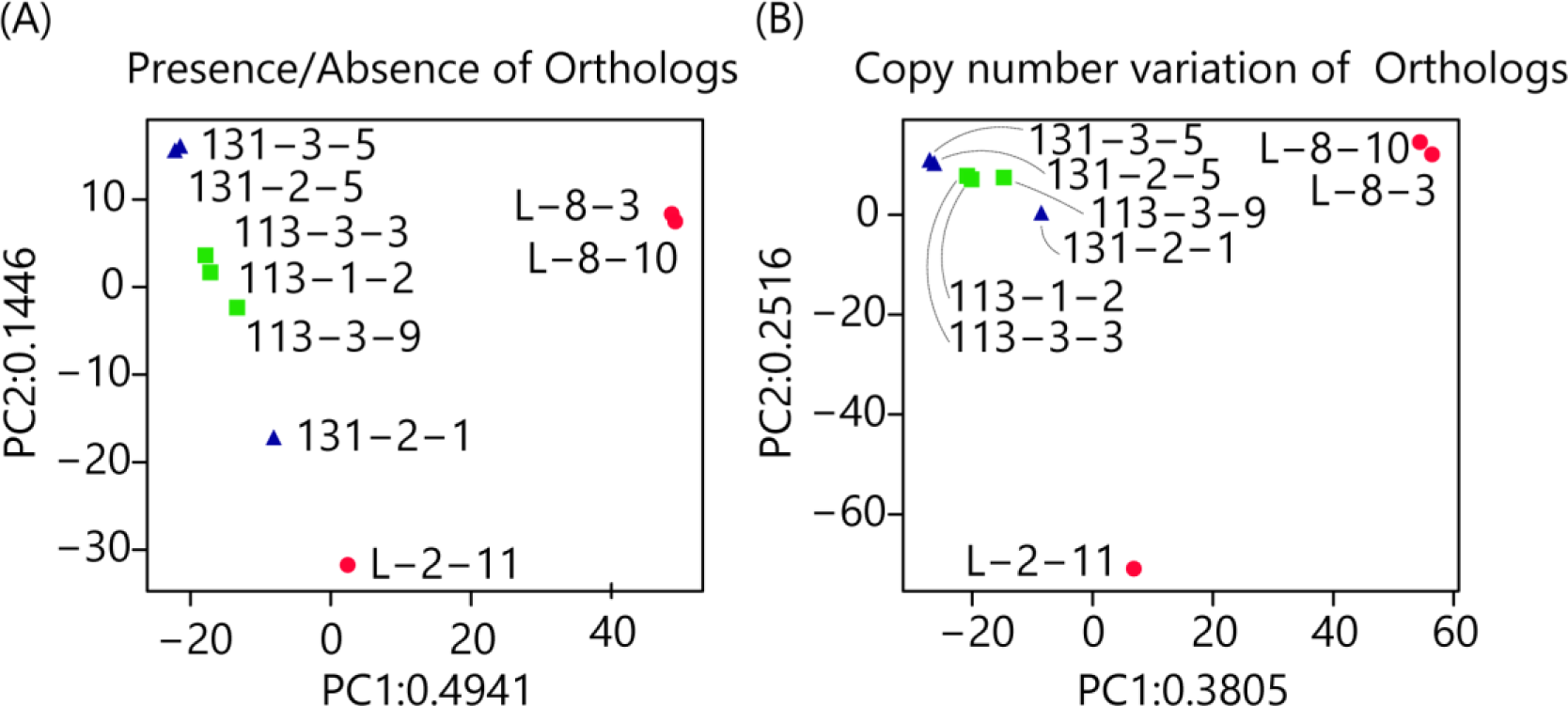
Principal component analysis (PCA) results of nine sequenced strains using ortholog profiles. (A) PCA based on the presence/absence of orthologs. (B) PCA based on copy number variations. Forms and colors of dots indicate the sampling localities of each rhizobial symbiont with the strain type designated in brackets: blue triangle, Aomori (131); green square, Tottori (113); and red circle, Miyakojima (L).

**Figure S2.**
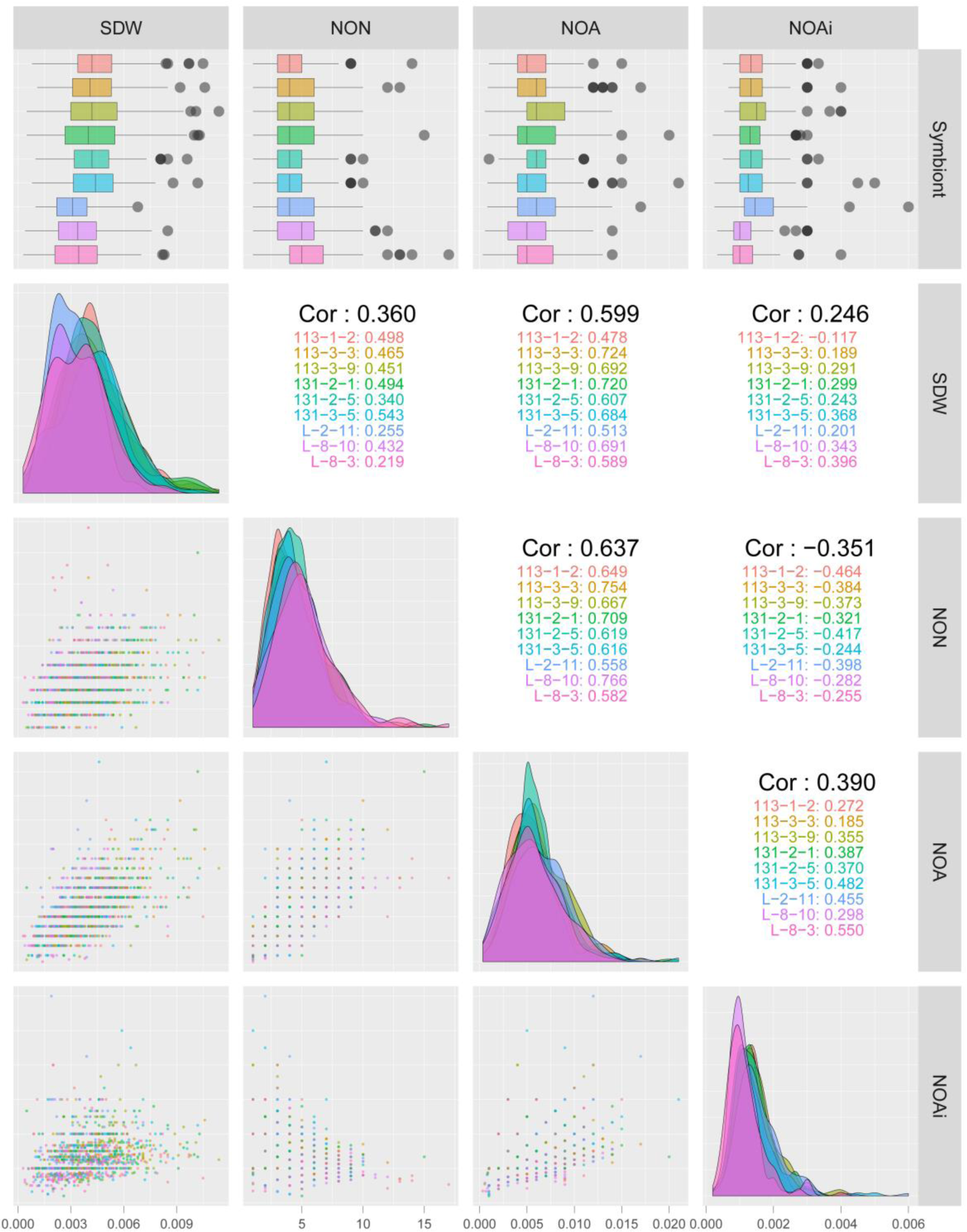
Correlations of phenotypes in rhizobial symbionts. Box plots show the distribution of phenotypic values for each rhizobial symbiont, and horizontal grey bars indicate the mean values. The x- and y-axes in each scatterplot represent phenotypic trait values: SDW, NON, NOA and NOAi. The diagonal plots are density curves for the individual points and Pearson’s correlation coefficients are given for all phenotypes (Cor; black). Each color indicates one rhizobial symbiont.

**Figure S3.**
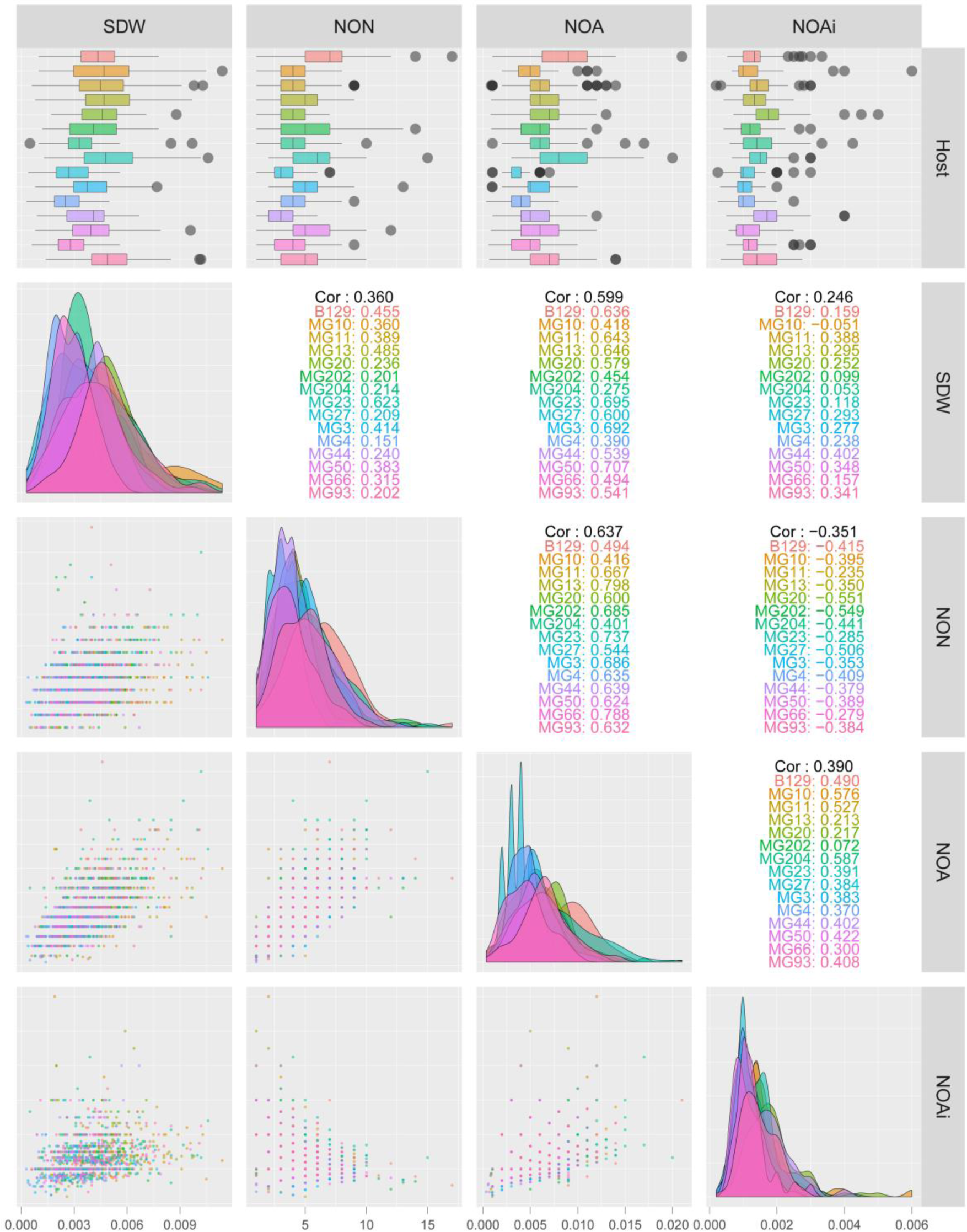
Correlations of phenotypes in *L. japonicus* accessions. Box plots show the distribution of phenotypic values for each accession, and horizontal grey bars indicate the mean values. The x- and y-axes in each scatterplot represent phenotypic trait values: SDW, NON, NOA and NOAi. The diagonal plots are density curves for the individual points and Pearson’s correlation coefficients are given for all phenotypes (Cor; black). Each color indicates one *L. japonicus* accession.

**Figure S4.**
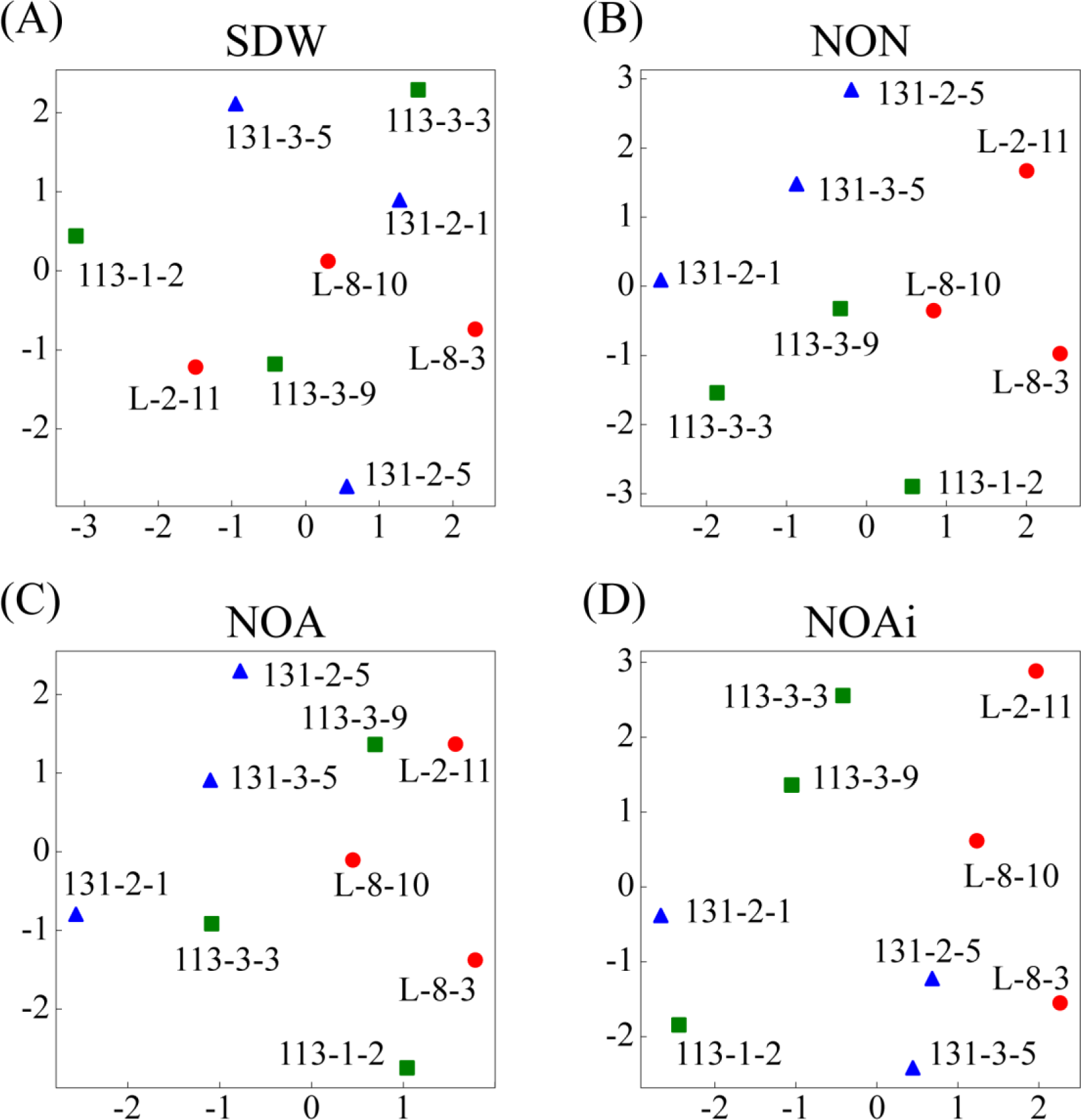
Non-metric multidimensional scaling of variations in G × G interactions of rhizobial symbionts: (A), SDW; (B), NON; (C), NOA; and (D), NOAi. Forms and colors of dots indicate sampling localities of each rhizobial symbiont with the strain type designated in brackets: blue triangle, Aomori (131); green square, Tottori (113); and red circle: Miyakojima (L).

**Figure S5.**
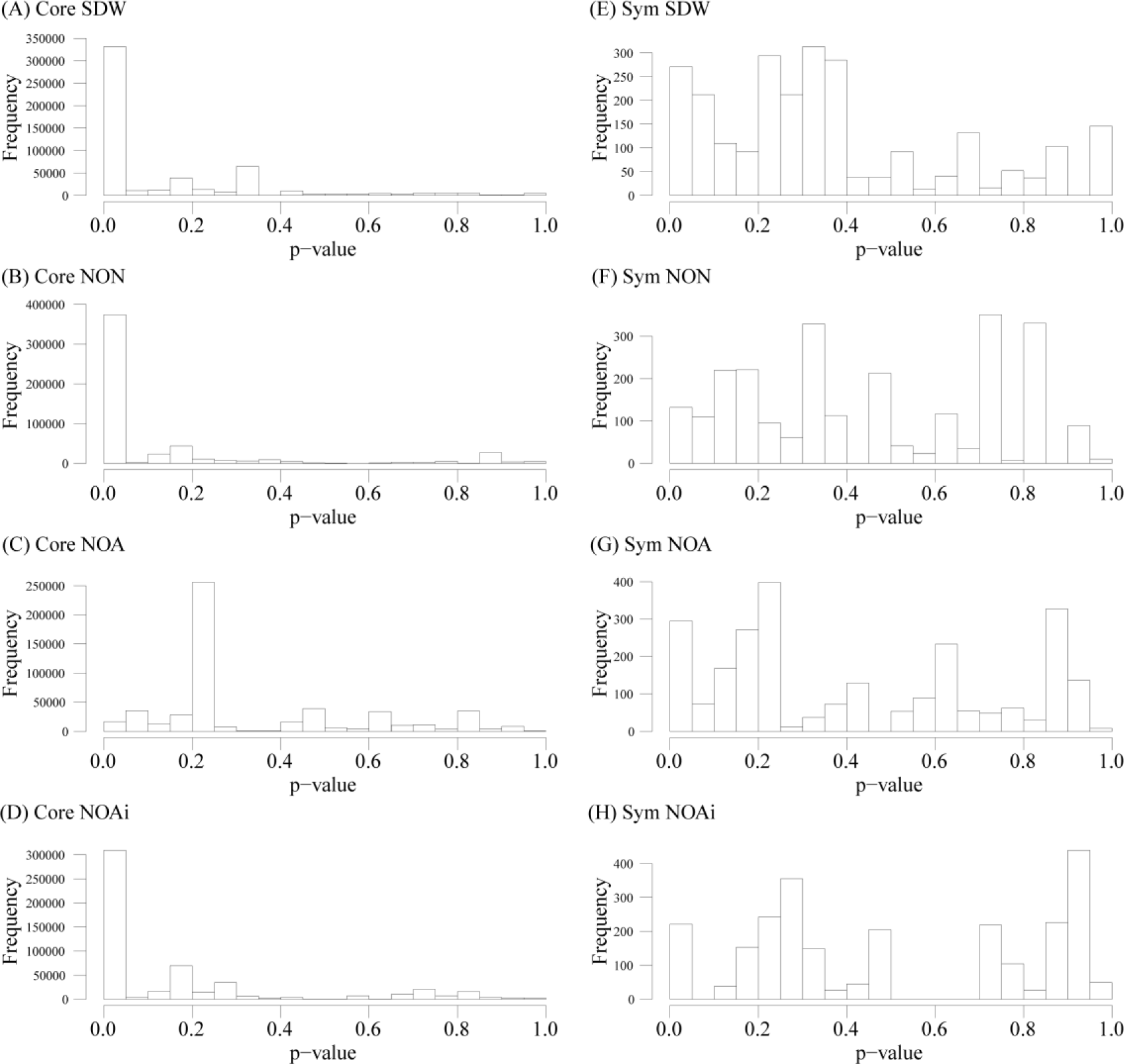
*P*-value distributions of correlations between variations in partner quality of each phenotype and the SNPs of the core genomes (A–D) and their symbiosis islands (E–H).

## SUPPLEMENTAL FILES

Supplemental Table 1: Summary of de novo assembly

Supplemental Table 2: List of host accessions

Supplemental Table 3: List of reference genomes

Supplemental Table 4: Summary of next-generation sequencing data

Supplemental Table 5: List of single copy orthologs

Supplemental Table 6: Results of TukeyHSD test

Supplemental Table 7: Results of ANOVAs 1

Supplemental Table 8: Results of ANOVAs 2

Supplemental Table 9: Results of Mantel test 1

Supplemental Table 10: Results of Mantel test 2

